# Biodiversity responses to forest cover loss: taxonomy and metrics matter

**DOI:** 10.1101/2023.05.21.541657

**Authors:** Clarissa de Araujo Martins, Olivier Pays, Franco L. Souza, Pierre-Cyril Renaud, Francisco Valente-Neto, Mauricio Silveira, Jose Ochoa-Quintero, Diogo B. Provete, Cyntia Cavalcante Santos, Isabel Melo, Marciel Elio Rodrigues, Samuel Duleba, André Valle Nunes, Oriana DJ. Ceballos-Rivera, Fabio de Oliveira Roque

## Abstract

The actions required for biodiversity conservation depend on species responses to habitat loss, which may be either neutral, linear, or non-linear. Here, we tested how taxonomic, functional, and phylogenetic diversity of aquatic insects, dragonflies, frogs, and terrestrial mammals, as well as their species composition respond to forest cover loss. We hypothesized that taxonomic, functional, and phylogenetic diversity would respond nonlinearly (thresholds) to forest cover loss. Our findings do not support the current idea that a single threshold value of forest cover loss is applicable across tropical regions, or that some biodiversity facets are consistently more sensitive than others across different taxa. Species compositional responses to forest cover loss showed general patterns with thresholds between 30-50%. These results highlight the importance to consider multiple biodiversity facets when assessing the effects of forest cover loss on biological communities.

## 1. Introduction

Human activities have substantially altered landscapes around the world, increasing species extinction rates (Dirzo et al., 2014). Tropical regions harbor most of the world’s biodiversity but are facing dramatic losses of forest cover. Therefore, these regions are losing biodiversity and ecosystem services at a high rate (Laurance et al., 2000; Barlow et al., 2018; Dalap-Corte et al., 2020). Forest cover loss may trigger different responses of biodiversity, including linear (proportional change), neutral (unrelated to habitat loss), or non-linear (disproportionate declines in biodiversity at specific habitat loss levels) relationships. If species respond non-linearly to habitat loss, biodiversity initially decreases proportionally to the forest cover remnant, until a particular level of forest loss is reached, after which an abrupt non-linear decline may occur (Andrén and Andren, 1994). In tropical agroecosystems, threshold levels usually range between 30 and 50% of forest cover, depending on the taxonomic group (Andrén and Andren, 1994; Pardini et al., 2010; Banks-Leite et al., 2014; Ochoa-Quintero et al., 2015; Melo et al., 2018; Arroyo-Rodríguez et al., 2020; Dala-Corte et al., 2020), however a recent overview of global evidence on the existence of forest cover thresholds in tropical regions and how to evaluate them highlighted large analytical and geographical bias (Shennan-Farpón et al., 2021).

Different mechanisms have been hypothesized to explain ecological thresholds. For example, closely related species share similar phenotypic traits and may respond synchronously to forest cover loss, causing drastic changes in community composition as disturbances exceed the tolerance levels of species (Dakos and Bascompte, 2014). Landscape percolation is another potential mechanism that can affect ecological thresholds and communities. Non-linear distance among habitat patches affects habitat permeability to movement by different species and taxonomic groups, eventually leading to isolation and local extinctions. The percolation associated with edge effects may have different consequences for organisms, including: (i) restricting species dispersal; (ii) changing abiotic conditions and resource availability; (iii) increasing the likelihood of ecological drift; (iv) changing ecological networks (Andrén and Andren, 1994; Lindenmayer et al., 2005; Morante-Filho et al., 2015; Matlaga, 2018; Boesing et al., 2018; Roque et al., 2018); or (v) causing cascading impacts on aquatic communities (Fuller et al., 2015; Raitif et al., 2019).

Most studies on biodiversity thresholds have focused on the effects of forest cover loss on taxonomic diversity metrics, such as species richness (Roque et al., 2018). However, biological diversity also includes alternative facets, such as variation among species in terms of their ecological functions and evolutionary relationships. Measuring the phylogenetic diversity of communities can explain the roles of species interactions and biogeographic histories in determining community structure and composition (Davies and Cadotte, 2016; Swenson, 2020), while functional diversity represents variability among species in terms of morphological, physiological, and ecological traits (de Bello et al., 2021). Although correlations among biodiversity facets have been reported along gradients of forest loss, exceptions have also been documented (Lindenmayer et al., 2015). For example, functional diversity may differ widely from phylogenetic diversity in situations where functional traits are under strong stabilizing selection or due to competitive interactions within lineages (Devictor et al., 2010; Cadotte et al., 2017; Tucker et al., 2018). Therefore, the evaluation of different biodiversity facets is important to obtain a more comprehensive understanding of the structure, composition, and dynamics of natural communities in response to forest cover loss (Devictor et al., 2010; Bovendorp et al., 2019).

Differences among taxonomic, functional, and phylogenetic diversity thresholds may be due to the sensitivity of facet responses to forest cover loss. Boesing et al. (2018) hypothesized that taxonomic diversity is the most sensitive of the three metrics, followed by functional diversity and finally phylogenetic diversity. They suggested that this pattern is because taxonomic diversity gives equal weight to each individual species with a range of tolerance or sensitivity levels, while functional diversity is less sensitive due to functional redundancy among species. Phylogenetic diversity is the least sensitive compared to the others because the loss of closely related species may have little influence on community phylogenetic diversity (Boesing et al., 2018). To date, this hypothesized threshold sensitivity sequence among biodiversity facets has been only supported by a study of Atlantic Forest birds. To obtain a more general understanding of how biodiversity facets respond to disturbances in human-modified landscapes, it is important to test Boesing et al. (2018) hypothesis for multiple taxonomic groups (Landeiro et al., 2018; Filgueiras et al., 2019).

Here, we tested the effects of forest cover loss on taxonomic, functional, and phylogenetic diversity in communities of aquatic insects, dragonflies, frogs, and terrestrial mammals. Our first hypothesis is that non-linear models, in comparison to linear and null models, would best explain the relationships between the percentage of forest cover and taxonomic, functional, and phylogenetic diversity. Due to the highly distinctive responses of species, our second hypothesis is that there would not be i) a single sequence on taxonomic, functional, and phylogenetic diversity thresholds and ii) universal response for multiple taxa, contrary to Boesing et al. (2018) hypothesis. Finally, we evaluated the variability in species composition for each taxonomic group along a gradient of forest cover loss. Our third hypothesis is that species composition would present points of abrupt change along the gradient of forest cover loss.

## 2. Materials and Methods

### 2.1 Study area

We compiled data from studies conducted on the Bodoquena plateau, Mato Grosso do Sul, central Brazil (Fig. 1). The vegetation consists mostly of deciduous and semideciduous forests, woodland, and arboreal savannas (*cerradão* and cerrado *stricto sensu*, respectively), and grasslands (Baptista-Maria et al., 2009). This region has one of the most extensive karst systems in Brazil and has become one of the most important ecotourist attractions in the world (Filho and Karmann, 2007). Economic activities in the region include cattle ranching, agriculture and, to a lesser extent, ecotourism. The Serra da Bodoquena National Park is a 77,000 ha Protected Area mainly consisting of pristine habitats. Surrounding the park, natural habitats have been widely converted into livestock pastures, corn, and soybean plantations, resulting in a landscape with forest patches.

**Figure 1.**
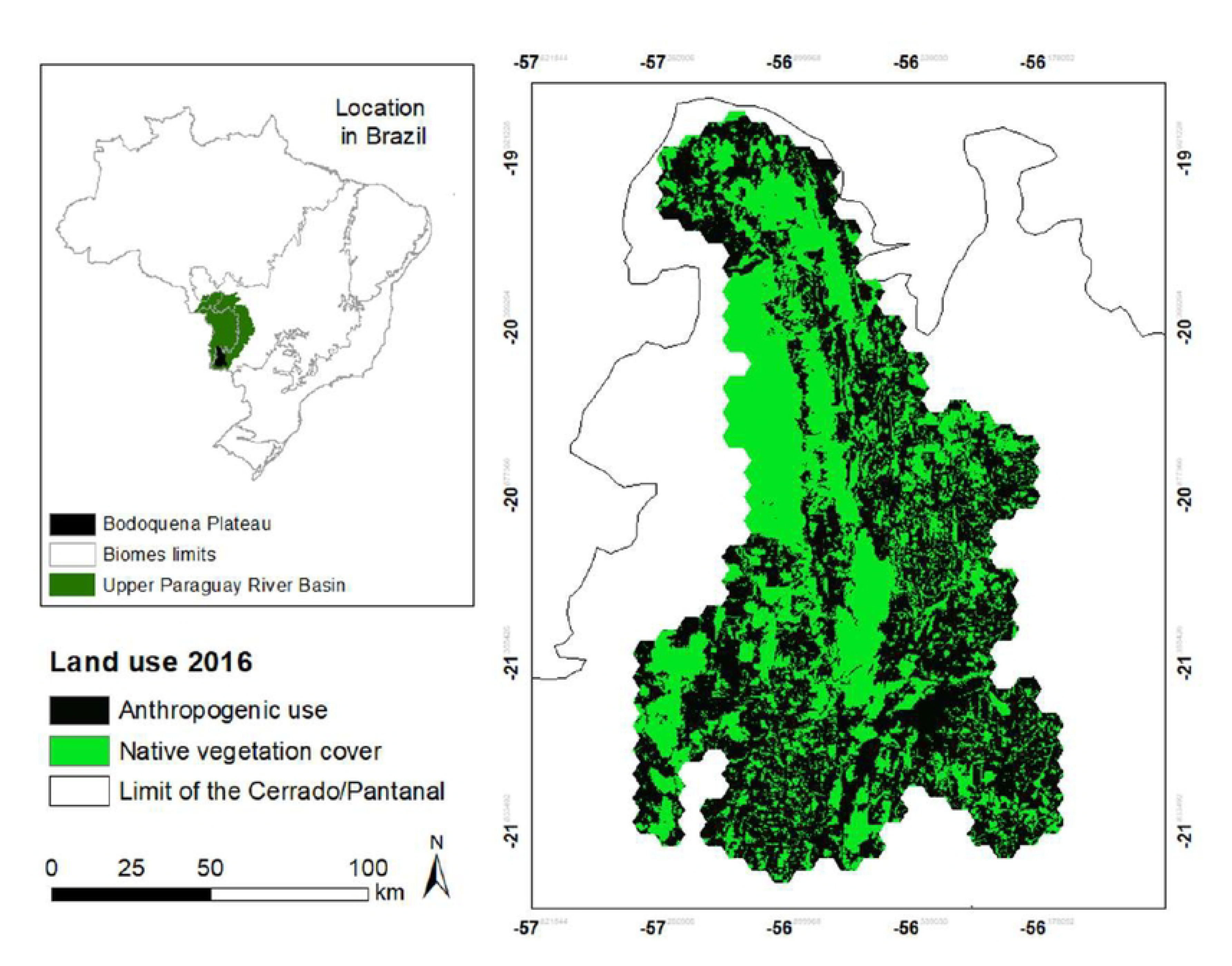
Map of study region, Bodoquena Plateau in Mato Grosso do Sul, state of Brazil. Geographic location and distribution of Brazilian biomes in the area.

### 2.2 Data collection and forest cover

Taxonomic groups (see sections below) were sampled at different spatial scale. We calculated forest cover associated with each sampling site by using different circular buffers surrounding the sampling point with the size of radius depending on the taxa (Galetti et al., 2013; Almeida-Gomes and Rocha, 2015; Rodrigues et al., 2016; Lashley et al., 2018). We considered forest cover as the area encompassed by deciduous and semideciduous forests and open-vegetated area according to MapBiomas. We selected the proportion of forest cover as our landscape metric, because it is one of the main variables that explains biodiversity distribution (Fahrig 2003; Quesnelle et al., 2013; Rodrigues et al., 2016). In addition, forest cover has commonly been used in studies of thresholds of species loss in different environments (Melo et al., 2018, Roque et al., 2019).

### 2.3 Aquatic insects

We used a dataset that includes samples from 46 stream sites collected in September 2013 (end of dry season) (Valente-Neto et al., 2018, Valente-Neto et al., 2020). At each site, aquatic insects were collected using a “D net” (0.5 mm mesh). Each sample consisted of 20 sweeps (sub-samples) of the upstream bed over an area of 1-meter length and 0.3-meter width. Thus, each sub-sample covered an area of 0.3 m^2^ of the stream bed and each sample covered an area of 6 m^2^ (Barbour et al., 2006). The sub-samples were distributed evenly along a 50-m stretch of the stream. The aquatic insects collected were preserved in a 4% formalin solution and then screened and identified mostly at genus level. For this dataset, the proportion of forest cover was calculated from a circular buffer (200 m radius) zone around each stream transect.

### 2.4 Odonates

For this dataset, we used a subset of samples from 98 stream sites collected by Rodrigues et al. (2016) for 1 hour along streams during sunny days between 10:00 and 15:00 h. At each site, a hand net was used to sample a 100-m linear transect. Sampling was conducted in streams with from 0 to 100% of native riparian vegetation remaining. To describe forest cover loss in the study area, we calculated the total proportion of riparian forest within a 250 m radius buffer zone that extended from the center of each sampled stream.

### 2.5 Anurans

We used a dataset that included 21 sampling sites in 15 forest fragments. Anurans were sampled using 1 h in 2 Km^2^ plots in each site. Sampling was performed by one observer at night (5 pm until midnight) using visual encounter (Crump and Scott Junior, 1994) and auditory survey (Zimmerman, 1994). We quantified forest cover within a 500-m buffer.

### 2.6 Terrestrial Mammals

For small (< 500 g body mass) terrestrial mammal sampling, we installed 26 Tomahawk ™ live traps on the ground and 26 Sherman traps in trees at 1.5 m along 1 Km transects. The distance between each pair of traps was about 20 m. Live traps were baited with fruits and pumpkins, and different mixtures of oats, peanut butter, and banana.

For medium to large (> 500 g body mass) terrestrial mammals, we recorded animals by actively searching along six 1-km transects in each of 15 previously described landscapes. One trained observer walked continuously along each transect from dawn to 11h. Species sightings were also recorded through opportunistic encounters. We also used camera-traps to record medium to large mammals over 25 days from June 2016 to December 2017 in each of the 15 landscapes. Each camera trap was set up following an identical standardized protocol to collect data on multiple mammal species. In total, 193 camera traps were installed in randomly chosen forested locations. We deployed a set of 10 to 15 cameras traps and adjusted the number of transect searches based on the percent of forest cover at each site, i.e., more cameras were used and more transects were conducted at sites with a higher percentage of forest cover. Camera trap delays for detecting passing animals were set at a frequency of 3 sec. We estimated species abundances by counting individuals recorded at least 3 minutes apart (Amiot et al., 2021).

To calculate the percentage of forest cover along transects and around camera trap sites within the 15 landscapes that represent a gradient from 0 to 100% of forest cover, we defined circular concentric buffers with a radius of 5 km for medium to large mammals and 1 km for small terrestrial mammals (Lashley et al., 2018).

### 2.7 Taxonomic Diversity

We quantified taxonomic diversity as the total number of species or taxa per sampling unit.

### 2.8 Functional Diversity

Response traits (*sensu* Violle et al., 2007, Luck et al., 2012, Moretti et al., 2017) were selected to reflect habitat use, body size, and species distribution area (Supplementary material, Table S1). These data were continuous, categorical (binary) or nominal (Supplementary material, Table S1), and were extracted from published literature and empirical data (Wilman et al., 2014, Amphibiaweb, 2019).

We used R v. 2.9.2 (R Core Team, 2009) to perform a Principal Coordinate Analysis (PCoA) with gower distance. We included only the first four axes in functional diversity analyses, because they captured a large proportion of the variation. We estimated functional diversity using the Rao quadratic entropy index. Rao’s QE is a measure of diversity that takes species differences (e.g., functional dissimilarity) into account (Pavoine et al., 2005; Ricotta and Szeidl, 2009; Mason et al., 2013), and incorporates species relative abundances. This index is a measure of functional divergence, since the higher the Rao QE, the greater the dissimilarity between species, and hence, the higher the functional divergence (Pena et al., 2017).

### 2.9 Phylogenetic Diversity

We quantified phylogenetic diversity using Faith’s PD, which is an index of phylogenetic richness (Tucker et al., 2017). For each taxonomic group, we used the total number of species sampled as the regional pool. Since PD increases monotonically with species richness, we also calculated the standardized effect size (SES.PD) (Tucker et al., 2017). Null communities were created by shuffling the tips of the phylogenetic tree 999 times (Kembel et al., 2010). Analysis was conducted in the R package ‘picante’ (Kembel et al., 2010).

We used specific phylogenetic tree hypotheses for each taxonomic group. Because there are no phylogenetic trees based on molecular data for aquatic insects and dragonflies, we used the phylogenetic tree based on topological data proposed by Saito et al. (2015, 2016). For anurans, we used the phylogenetic tree proposed by Pyron and Wiens 2011, and for terrestrial mammals, we used the phylogenetic tree proposed by Bininda-Emonds et al. (2007).

### 2.10 Data analysis

To test the first (non-linear models best explain metric communities) and second hypotheses (no multiple taxa threshold) we built, for each biodiversity metric, a null model, a generalized linear model (GLM), and a piecewise analysis. The null model tested the absence of effect of forest cover on biodiversity metrics and is suitable to verify if GLM and piecewise analysis were better than expected by chance. The GLM assumes a linear relationship between predictor (vegetation cover) and response variables (taxonomic, functional, and phylogenetic diversity metrics). For the GLMs, we used a Poisson distribution with log link for taxonomic diversity, and Gaussian distribution for functional and phylogenetic diversities.

For the piecewise regression, we used segmented regression analysis that assumes breakpoint indicating thresholds caused by abrupt losses of diversity (taxonomic, functional, or phylogenetic) (Muggeo et al., 2014). Finally, to identify the best-fit models for each biodiversity metric, we compared models using the Akaike Information Criterion corrected for small sample sizes (AICc). Analyses were conducted using the R packages "lme4" (Bates et al., 2015), "lmerTest" (Kuznetsova et al., 2017) and "segmented" (Muggeo et al., 2014).

To test our third hypothesis (species composition responds strongly to a gradient of forest cover), we used the Threshold Indicator Taxa Analysis “TITAN” (Baker and King, 2010). TITAN identifies threshold(s) based on the point of change along environmental gradients for each taxon. To carry out the TITAN analysis, we only used taxa with more than five records. We log-transformed species abundances to reduce the influence of variable taxa on the calculations of indicator scores in each dataset (1,000 repetitions and 1,000 bootstraps). We conducted analyses in the R v. 2.9.2 (R Core Team, 2009) package “TITAN” (Baker and King, 2010).

## 3. Results

The responses of biodiversity facets to the gradient of forest cover loss were highly variable among taxonomic groups. A linear model provided the best fit for medium to large terrestrial mammal taxonomic diversity and the relationship between taxonomic diversity and percentage of forest cover was positive (Fig. 2). For anuran taxonomic diversity, the piecewise model provided the best fit and it detected a threshold at around 59% of forest cover. Intermediate levels of forest cover had the lowest anuran taxonomic diversity (Fig. 2; Supplementary material, Table S2). For aquatic insects and odonates, the null model provided the best fit (Supplementary material, Table S2).

**Figure 2.**
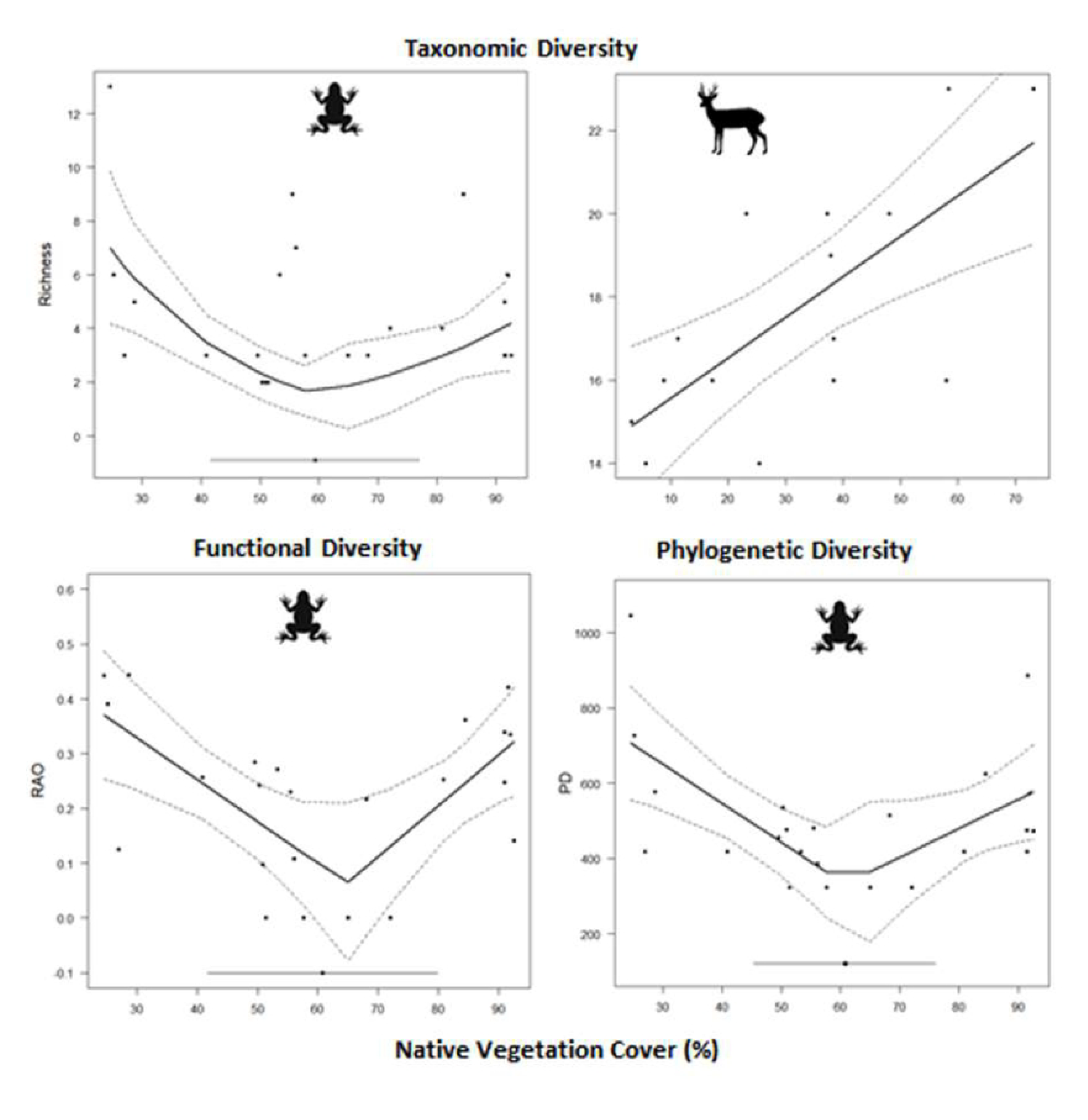
Relationships between three categories of biodiversity metrics (i.e., taxonomic, functional, and phylogenetic) describing different taxonomic groups and percentage of forest cover. The solid line represents the logarithmic trend line, the dashed lines represent standard error and the circles on straight lines at the lower portions of some graphs represent thresholds. The top 2 graphs show taxonomic diversity trends for anurans (left) and medium to large mammals (right), while the middle and bottom graphs show the functional diversity and phylogenetic diversity trends, respectively, for anurans.

The piecewise model provided the best fit for anuran functional diversity, and it showed a threshold at 65% forest cover. Anurans had their lowest functional diversity between 55-70% of forest cover (Fig. 2; Supplementary material, Table S2). The piecewise model provided the best fit for anuran phylogenetic diversity and had a threshold of around 62% forest cover. The anuran phylogenetic diversity pattern was quite like that of anuran taxonomic and functional diversity (Fig. 2). The highest values of phylogenetic diversity were observed at high or low percentages of forest cover (Fig. 2; Supplementary material, Table S2).

Species composition showed thresholds between 30 and 60% of forest cover for three of the four taxonomic groups (Fig. 3). Aquatic insects showed a community composition threshold around 40% of forest cover, while odonates had a threshold close to 45% (Fig. 3). For medium to large terrestrial mammals, we detected thresholds around 35% of forest cover (Fig. 3). Results for anurans and small mammals did not show species with adequate reliability index values, so we were unable to obtain meaningful results from species compositional analyses. Reliable indicators of forest cover change included six aquatic insect species, 14 odonate species, and 6 mediums to large terrestrial mammal species (Supplementary material, Figure S1, Table S3).

**Figure 3.**
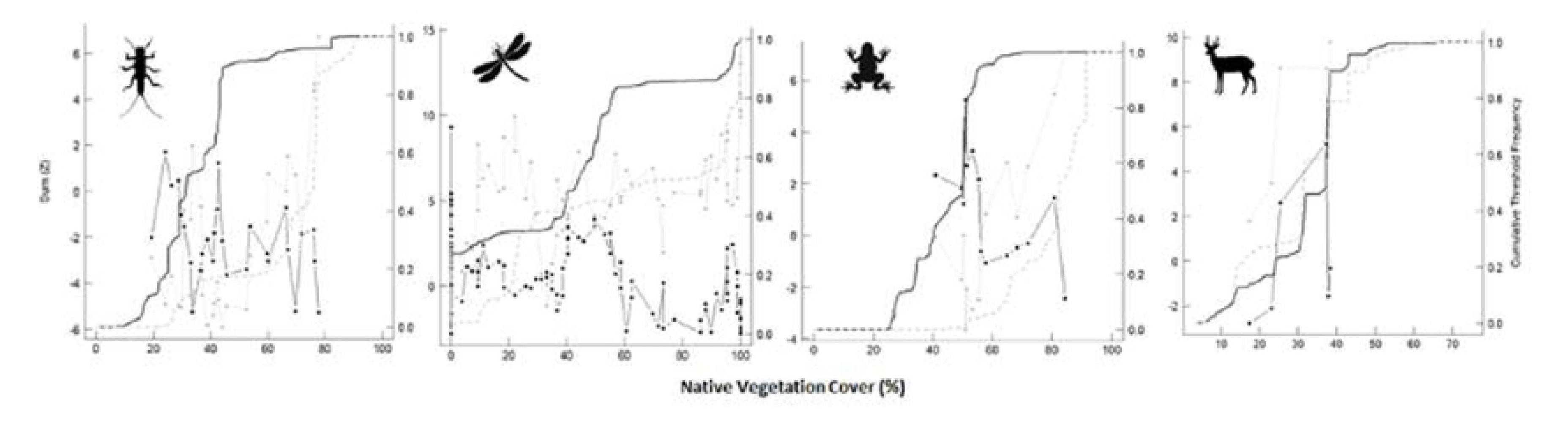
Results of TITAN analyses showing sum (Z -) and sum (Z +) values corresponding to all candidates change points along a forest cover environmental gradient. Black and gray lines without point symbols represent the cumulative frequency distributions of change points for 500 bootstrap replicates for sum (Z -) and sum (Z +), respectively. Steep sections of the distributions represent community composition thresholds.

## 4. Discussion

A growing number of studies from tropical regions have addressed the idea that biodiversity responses to forest cover loss are non-linear phenomena (Pardini et al., 2010; Banks-Leite et al., 2014; Rodrigues et al., 2016; Arroyo-Rodríguez et al., 2020), but large gaps of knowledge remains (Shennan-Farpón et al., 2021). These studies suggested that some taxonomic groups show clear disproportional changes – or threshold – caused by small change in forest cover loss (Banks-Leite et al., 2014; Boesing et al., 2018; Arroyo-Rodríguez et al., 2020). Also, Boesing et al. (2018) suggested a predictable sequence of responses considering different biodiversity components. One of the implications of these studies is a proposal to define a “generalizable ecological threshold value”. This value means a given forest loss percentage that can be used for setting regulatory limits on forest conversion and for defining conservation actions involving societal choices and negotiations over values and aims (Banks-Leite et al., 2014; Arroyo-Rodríguez et al., 2020; Dala-Corte et al., 2020; Hillebrand et al., 2020). Our study does not support the use of a generalizable ecological threshold value, because different components of biodiversity showed varied responses to forest cover loss. However, this does not mean that the idea of using ecological thresholds to orient conservation and restoration strategies should be discarded. Instead, our findings suggest that these efforts will be most effective when selecting sensitive metrics and best fit models to conservation problems that require knowledge of multi-taxa taxonomic, functional, phylogenetic and community compositional responses to vegetation loss. Adapting these models for specific taxonomic groups and regions will be a critical step in selecting sensitive bioindicators of the effects of forest loss, and based on our results, non-linear responses should be considered during the process of model development.

Following previous works (e.g., Banks-Leite et al., 2014; Boesing et al., 2018), we expected to find nonlinear relationships between taxonomic, functional, and phylogenetic diversity and forest cover. However, our results showed idiosyncratic responses, including linear, non-linear, and null relationships between metrics of different taxonomic groups and the vegetation cover gradient. Consequently, taxonomic, functional, and phylogenetic diversity did not show threshold values or response patterns that can be generalized across multiple aquatic and terrestrial taxa, showing context-dependent responses of metrics and taxonomic groups (Lindenmayer et al., 2005; Roque et al., 2018; Dala-Corte et al., 2020). Only when we focused on community compositional changes did a more general pattern emerge, showing thresholds between 30-60% of forest cover for some taxonomic groups.

One possible explanation for the lack of similar threshold responses among taxonomic groups and the unpredictable sequence of different biodiversity metrics responses to forest cover loss (Boesing et al., 2018) is that some environmental gradients, such as forest cover, may not be associated with effects that are strong enough to elicit similar responses from multiple taxa with variable environmental tolerances (Heino et al., 2008; Hidasi-Neto et al., 2019). For example, some altered landscapes provide a diversity of habitats that support multiple species with different traits and habitat requirements. Another potential explanation is “historical filtering”, which may explain some of the results from our study. In the Bodoquena Plateau region, vegetation experienced cycles of forest contraction and expansion during the quaternary (Silva et al., 2006; Cáceres et al., 2007). As such, species may already be adapted to variable percentages of forest cover, resulting in resilient communities that are, consequently, less sensitive to forest cover loss (Melo et al., 2018). Indeed, many lineages from different groups, very common in the Bodoquena Plateau, such as microhylid frogs, zygopterans (Odonata) and xenarthrans (Mammalia) use both open and forested vegetation formations (Almeida-Gomes and Rocha, 2015; Fischer et al., 2018). Furthermore, the idiosyncratic responses of multiple taxa to forest cover loss may be explained by ecological compensatory dynamics. For example, as forest cover is reduced, the populations of some species may decline while others increase, causing species turnovers that are masked by aggregate community metrics, like species richness, that remain relatively constant as vegetation cover declines. However, it is important to note that for medium to large terrestrial mammals we found linear loss of taxonomic diversity as forest cover declined.

Overall, the responses of taxonomic, functional, and phylogenetic diversity to forest cover loss showed little evidence of ecological thresholds. Our results showed that only anurans responded non-linearly to forest cover loss, with a threshold around 60%. Additionally, anuran taxonomic, functional, and phylogenetic diversities showed a similar V-shaped relationship with forest cover. These patterns were surprising, because evidence suggested that amphibians are an environmentally sensitive group and show clear ecological thresholds along gradients of vegetation loss (Roque et al., 2018). The V-shaped relationship we observed for anurans may be explained by the turnover of species along the gradient of forest cover loss, which also occurred at the phylogenetic level. As forest cover decreased, some clades were replaced by others with greater tolerance to forest cover loss. On the Bodoquena Plateau, even landscapes with lower levels of forest cover may have water supply, such as artificial aquatic habitats for cattle management in pastures that allow the occurrence of anuran lineages adapted to open areas.

In contrast to the biodiversity metrics, species composition showed clear threshold patterns to forest cover loss for most groups (i.e., aquatic insects, dragonflies, and medium to large mammals). This may indicate that aggregate measures, like metrics of taxonomic, functional, and phylogenetic diversity, provide incomplete information for understanding the response of different taxonomic groups to forest cover loss, and that information about individual species may be critical to the selection of the most sensitive bioindicators of forest cover change (Lindenmayer et al., 2005). Our results for composition showed thresholds between 30-60% and high variability around an average of 45% of forest cover. This high variability may be related to specific requirements and dependencies of species to forest cover. Aquatic insects and terrestrial mammals had sharper thresholds (around 38%), probably because some species are highly dependent on forest resources and environmental conditions (Valente-Neto et al., 2021). For example, some species require fruits, leaves, wood or shady conditions to survive (Valente-Neto et al., 2018). Furthermore, forest cover loss can increase sediment inputs and luminosity in aquatic systems, causing synchronous losses of forest-dependent aquatic species (Raitif et al. 2019; Brito et al., 2020, Valente-Neto et al., 2021). Interestingly, we found similar threshold values for terrestrial and aquatic communities, leaving open the potential for development of negative feedback cycles involving both terrestrial and aquatic ecosystems as vegetation losses exceed 40%. Taken together, the results for aquatic insects, dragonflies and medium to large mammals partially support the idea that ecological thresholds occur around 40% for tropical regions, as proposed by Andrén and Andren (1994), Pardini et al. (2010) and Melo et al. (2018). However, as recently highlighted for aquatic systems (Dala-Corte et al., 2020), a single threshold value should not be applied as a panacea. For example, the threshold we observed for anurans was around 60% of forest cover. So, applying a 40% threshold to set forest conversion limits would increase the chances for abrupt disproportionate losses of anuran species.

Our findings also have some practical implications. First, we suggest avoiding the use of aggregate metrics (e.g., taxonomic diversity) to set regulatory limits and define conservation actions based on ecological thresholds. As an alternative, we suggest that regulations and actions be based on the threshold patterns observed for a set of the most sensitive species or groups considering different biodiversity dimensions. This will increase the effectiveness of monitoring programs to detect impacts, in addition to saving time and funds. Secondly, developing management actions and policies at the landscape scale (e.g., regulatory limits for forest cover) based on ecological threshold patterns is a promising avenue that can inform conservation planning (Banks-Leite et al., 2012, Arroyo-Rodríguez et al., 2020). However, it will be important to consider regional contexts and the unique responses of multiple taxa and metrics, rather than relying on a general threshold value that is appropriate for a specific group, but potentially harmful to other components of biodiversity.

## Acknowledgments

This study was financed in part by the Coordenação de Aperfeiçoamento de Pessoal de Nível Superior – Brasil (CAPES) – Finance Code 001. The Project also received financial support from Fundação de Amparo à Pesquisa de Mato Grosso do Sul (Fundect), and Conselho Nacional de Desenvolvimento Científico e Tecnológico (CNPq). FVN is supported by grant 2021/13299-0, São Paulo Research Foundation (FAPESP).

## Author Contributions

CAM, FOR conceived the idea. CAM, OP, MS and DBP developed the statistical analysis. FVN, CCS, IM, MER, SD collected the data. CAM analyzed the data. CAM and FOR wrote the manuscript. FLS, PCR, FVN, AVN, JOQ and OCR drafted or revised the manuscript. All authors revised the manuscript or provided editorial advice.

## Conflicts of interest

We have no competing interests.

## Ethics approval

Not applicable.

## Consent to participate

Not applicable.

## Consent for publication

Not applicable.

## Availability of data and material

All data are included on Supplementary Material.

## Code availability

Not applicable.

## Notes

### Competing Interest Statement

The authors have declared no competing interest.

